# Metagenomic analysis of milk microbiota in the bovine subclinical mastitis

**DOI:** 10.1101/2023.05.09.539964

**Authors:** Giulia Alessandri, Elena Sangalli, Mario Facchi, Federico Fontana, Leonardo Mancabelli, Gaetano Donofrio, Marco Ventura

## Abstract

Subclinical mastitis is one of the most widespread diseases affecting dairy herds with detrimental effects on animal health as well as on milk productivity and quality. Despite the multi-factorial nature of this intramammary infection, the presence of pathogenic bacteria is regarded one of the main drivers of subclinical mastitis, leading to a disruption of the homeostasis of the bovine milk microbial community. However, the bovine milk microbiota alterations associated with subclinical mastitis still represents a largely unexplored research area. In this context, the species-level milk microbiota of a total of 75 milk samples, collected from both healthy and subclinical mastitis-affected cows from two different stables, was deeply profiled through an ITS, rather than a traditional, and less informative, 16S rRNA gene microbial profiling-based sequencing. Surprisingly, the obtained data of the present pilot study, not only revealed that subclinical mastitis is characterized by a reduced number of species in the bovine milk microbiota, but also that this disease does not induce standard alterations of the milk microbial community across stables. In addition, a flow cytometry-based total bacterial cell enumeration highlighted that subclinical mastitis is accompanied by a significant increment in the number of milk microbial cells. Furthermore, the combination of the metagenomic approach and total bacterial cell enumeration allowed to identify different potential microbial marker strictly correlated with subclinical mastitis across stables.

## Introduction

Bovine mastitis is a worldwide recognized disease affecting dairy cows with devastating impacts on productivity, milk quality, and animal well-being (1-3). Clinically defined as an inflammation of the mammary gland, bovine mastitis is caused by multi-etiological agents, including several microbial and environmental predisposing factors (3-5). Based on the severity of the symptoms, this disease is classified into clinical or subclinical mastitis (SM), both accompanied by high milk somatic cell count (4, 5). However, if the former is distinguished by evident physiological alterations, including swelling and inflammation of the mammary gland as well as changes in milk color, consistency, and yield, the latter is characterized by a shortage of visible clinical symptoms, yet a damage in lactation performance, immune function, and alteration of the normal metabolic activities (3, 5-7). Consequently, due to the long latency period and the lack of obvious clinical signs that prevent prompt interventions to limit its spread, SM incidence is significantly higher than that of clinical mastitis, accounting for approximately 90% of bovine mastitis cases (7). Furthermore, despite its multi-factorial nature, SM generally occurs as a result of intramammary infection induced by specific pathogenic bacteria that not only trigger inflammation, leading to detrimental effects for both mammary tissue and bovine physiology, but also disrupt the homeostasis of the bovine milk microbial community with a consequent overgrowth of these pathogenic microorganisms and potential risk of their transmission to healthy cows (5, 8-10).

However, despite the relevant role played by bacteria in SM etiology, the milk microbial composition associated to this clinical status is still far from being completely dissected. Indeed, most of publicly available metagenomic studies only employed 16S rRNA gene microbial profiling-based sequencing, thus preventing an accurate and complete characterization of the bovine milk microbiota associated to SM down to the species level (3, 6, 7, 11, 12). At the same time, studies limited to culture-dependent investigations, despite being able to identify the presence of underrepresented pathogenic microorganisms in subclinical bovine milk whose detection can escape metagenomics due to the intrinsic limit of this molecular approach (13), do not allow to obtain an accurate overview of how the milk microbiota can change during SM (14-16).

In this context, to evaluate possible species-level alterations of bovine milk microbial composition due to SM, a total of 72 milk samples, subdivided into 38 and 34 milk samples from healthy and SM-affected cows, respectively, were collected from two different stables. Subsequently, samples were simultaneously subjected to an Internally Transcribed Spacer (ITS) microbial profiling sequencing and to a flow cytometry-based total bacterial cell enumeration. The analysis of the microbial profiles revealed that environmental factors play a crucial role in modulating the taxonomic composition of milk microbiota and, therefore, to avoid biases related to environmental factors, samples were analyzed separately based on their stable of origin. In this context, the comparison of milk microbial community between healthy and diseased cows from the two stables highlighted that SM does not induce unique alterations in the bovine milk microbiota, but rather, the microbial modulation seems to be stable-dependent. In support of this finding, diverse bacterial species have been identified to be associated to SM, and therefore as microbial marker closely associated with subclinical mastitis, including *Corynebacterium bovis, Corynebacterium xerosis*, and *Streptococcus uberis*, between the two considered stables. Furthermore, total bacterial cell enumeration highlighted that SM is strictly associated with a significant increment of the total microbial cells present in the milk samples.

## Experimental Procedures

### Ethical statement

All the dairy cows involved in this study were reared in commercial private farms and were not subjected to any invasive procedures. Milk samples used for the analyses were collected during the daily milking procedure in according to the International Committee for Animal Recording procedures (ICAR https://www.icar.org/index.php/icar-recording-guidelines/).

### Sample collection and clinical health status screening

Raw milk samples were collected from a total of 72 dairy cows, divided into 38 healthy cows and 34 cows affected by SM, from two different farms located in the North of Italy (Table S1). Per each cow, two milk samples were sterilely collected by hand from all milking quarters during the morning milking. One of the two milk samples of each milking quarter was collected in bronopol tubes for Somatic Cell Count (SCC) analysis. Before collection, the teat-ends were cleaned and properly disinfected with 70% ethanol, while the first milk jets were discarded. Furthermore, only milk samples from dairy herds that had not undergone any antibiotic treatment during the two months prior sample collection were included in this study. Once collected, milk samples were refrigerated and immediately shipped to the laboratory where 50 ml were preserved at -20°C for DNA extraction and flow cytometry-based cell enumeration, while the other 50 ml in bronopol tubes were stored at 4°C for SCC analysis. The latter was performed by using the XX. the cut-off value set for the determination of SM was SCC > 200,000 cells/ml, as previously described (16, 17).

### DNA extraction and microbial ITS profiling

Raw milk samples were subjected to DNA extraction using the DNeasy PowerFood Microbial Kit (Qiagen, Germany), following the manufacturer’s instructions. Subsequently, the Internal Transcribed Spacer (ITS) sequences were amplified from extracted DNA using the primer pair UNI_ITS_fw (5′-KRGGRYKAAGTCGTAACAAG-3′) and UNI_ITS_rv (5′-TTTTCRYCTTTCCCTCACGG-3′), targeting the entire spacer region between the 16S rRNA and 23 rRNA genes within the rRNA locus, as previously described (18). Illumina adapter overhang nucleotide sequences were added to the ITS amplicons, which were further processed using the 16S Metagenomic Sequencing Library Preparation Protocol (Part No. 15044223 Rev. B – Illumina). Amplifications were carried out using a Verity Thermocycler (Applied Biosystem, USA). The integrity of the PCR amplicons was analyzed by gel electrophoresis. DNA products obtained following PCR-mediated amplification of the ITS region sequences were purified by a magnetic purification step employing the Agencourt AMPure XP DNA purification beads (Beckman Coulter Genomics GmbH, Brea, USA), to remove primer dimers. DNA concentration of the amplified sequence library was determined by a fluorometric Qubit quantification system (Life Technologies, USA). Amplicons were diluted to a final concentration of 4 nM, and 5 μl of each diluted DNA amplicon sample were mixed to prepare the pooled final library. Sequencing was performed using an Illumina MiSeq sequencer with MiSeq reagent kit v3 chemicals, using 300 cycles.

### ITS microbial profiling analysis

After sequencing, the obtained .fastq files were processed using the METAnnotatorX2 pipeline (19). Specifically, paired-end reads were merged, and quality control retained only sequences with a minimum length of 100 bp and a mean sequence quality score of >20. Sequences with mismatched forward and/or reverse primers were omitted. Furthermore, sequences were filtered to remove *Bos taurus* DNA.

### Evaluation of bacterial cell density by flow cytometry

For total bacterial cell count, each milk sample was 10,000 diluted in physiological solution (Phosphate Buffered Saline, PBS, pH 6.5). Subsequently, 1 ml of the obtained bacterial cell suspension was stained with 1 μl of SYBR Green I (Invitrogen, Waltham, USA) (1:100 diluted in dimethyl sulfoxide), vortex-mixed, and incubated in the dark for at least 15 min before measurement. All count experiments were performed using an Attune NxT flow cytometry (ThermoFisher Scientific, Waltham, USA) equipped with a blue laser set at 50 mV and tuned at an excitation wavelength of 488 nm. Multiparametric analyses were performed on both scattering signals, i.e., side scatter and forward scatter, while SYBR Green I fluorescence was detected on the BL1 530/30 nm optical detector. Cell debris was excluded from acquisition analysis by setting a BL1 threshold. In addition, to exclude remaining background events and obtain an accurate microbial cell count, the gated fluorescence events were evaluated on the forward-sideways density plot, as previously described (20). All data were statistically analyzed with the Attune NxT flow cytometry software.

### Statistical analyses

Eigenvalue scores were retrieved from a Bray-Curtis dissimilarity matrix based on the taxonomical profiles of samples. Two-dimensional PCoA representation of eigenvalue scores were carried out using OriginLabPro 2021b. Ellipses in the PCoA were drawn based on standard deviation of each group. The confidence limit for ellipses was set to 0.95. PERMANOVA statistical analyses were performed using Rstudio software. Furthermore, SPSS software was used to compute the independent Student’s T-test statistical analyses.

### Data availability statement

ITS microbial profiling data were deposited in the NCBI-related SRA database with the accession number PRJNA942519.

## Results and Discussion

### Characterization of the microbial community of milk samples from healthy and subclinical mastitis-affected cows

To highlight possible species-level taxonomical differences in the bovine milk microbial community between healthy cows and cows affected by SM, a total of 72 milk samples were collected, divided into 38 milk samples from healthy cattle and 34 milk samples from cows with subclinical mastitis (Table S1). Subsequently, the microbial DNA extracted from each milk sample was subjected to an ITS microbial profiling, as previously described (18). Illumina sequencing generated a total of 4,342,880 reads with an average of 60,317 reads per sample, reduced to a total of 2,085,812 reads with an average of 28,969 reads per sample after filtering for quality and *Bos taurus* DNA (Table S2).

The species richness analysis revealed that the number of bacterial species present in healthy cow milk is significantly higher than that of the milk collected from cows with SM, with an average number of species of 54 and 35, respectively (Student’s T-test p-value < 0.01) (Figure S1). Thus, suggesting that subclinical mastitis is characterized by a significant reduction of milk microbial biodiversity, a condition that is frequently encountered in microbial communities associated with various diseases (32864871, 35038617). However, a Bray-Curtis dissimilarity-based beta-diversity analysis, represented through a Principal Coordinate Analysis (PCoA), revealed that environmental factors (R^2^ = 0.181 and PERMANOVA *p*-value = 0.001), i.e., the different stable from which samples were collected, seemed to have a higher impact on the modulation of milk microbial biodiversity than cow clinical status (R^2^ = 0.038 and PERMANOVA *p*-value = 0.003), with a clear separation of samples according to their stable of origin (Figure S1). Thus, indicating that different diets, environments, and litters could play a crucial role in the modulation of the bovine milk microbiota regardless of the cow clinical status. Furthermore, the PCoA showed that two samples from stable 1 displayed a microbial taxonomic profile that strongly differed from that of the other samples from the same stable (Figure S1). Therefore, they were considered as outliers and eliminated from subsequent analysis.

### Stable-related differences in the taxonomic composition of milk samples between healthy and subclinical mastitis-affected cows

Based on the above findings according to which the exposure to different diet, litters, and breeding management, strongly influenced the milk microbial communities, to avoid biases related to environmental factors, the collected samples were separately analyzed according to their stable of origin to evaluate possible differences in the taxonomic composition of milk samples from healthy and SM-affected cows. A separation that was possible because, despite the small number of sampled stables, the number of milk samples is balanced between healthy and subclinical mastitis-affected cows within each stable (Table S1). In this context, the species richness analysis highlighted that only for one of the two stables, i.e., stable 2, the number of bacterial species was significantly higher in the milk samples from healthy cows when compared to that from cows with SM (Student’s t-test *p*-value < 0.001), counting an average number of microbial species of 55 and 29, respectively (Figure 1 and Table S3). However, even if not statistically significant (Student’s T-test *p*-value = 0.482), a slight increase in the average number of microbial species was observed in healthy cow milk samples from stable 1 respect to the SM-affected cows, passing from an average of 54 to 50 bacterial species, respectively (Figure 1 and Table S3). Thus, strengthening the abovementioned notion that, even if not always statistically significant, SM is characterized by a general reduction of milk microbial biodiversity. In addition, in-depth insights into the microbial biodiversity of milk samples divided per stable and represented through a PCoA highlighted that the SM played a significant role (R^2^ = 0.0894 and PERMANOVA *p*-value < 0.001) in the modulation of the taxonomic composition of milk samples from stable 2 with a clear separation of samples according to their clinical status, while the microbial communities of milk samples from stable 1 did not differ in biodiversity between healthy and SM cows (R^2^ = 0.024 and PERMANOVA *p*-value = 0.798) (Figure 1). Thus, suggesting that subclinical mastitis does not always induce a drastic modulation in the taxonomic composition of the milk microbial communities. Conversely, this finding indicates that, depending on the environmental factors, SM is characterized by a different alteration of the bovine milk-related microbial community biodiversity and species richness.

**Figure 1:**
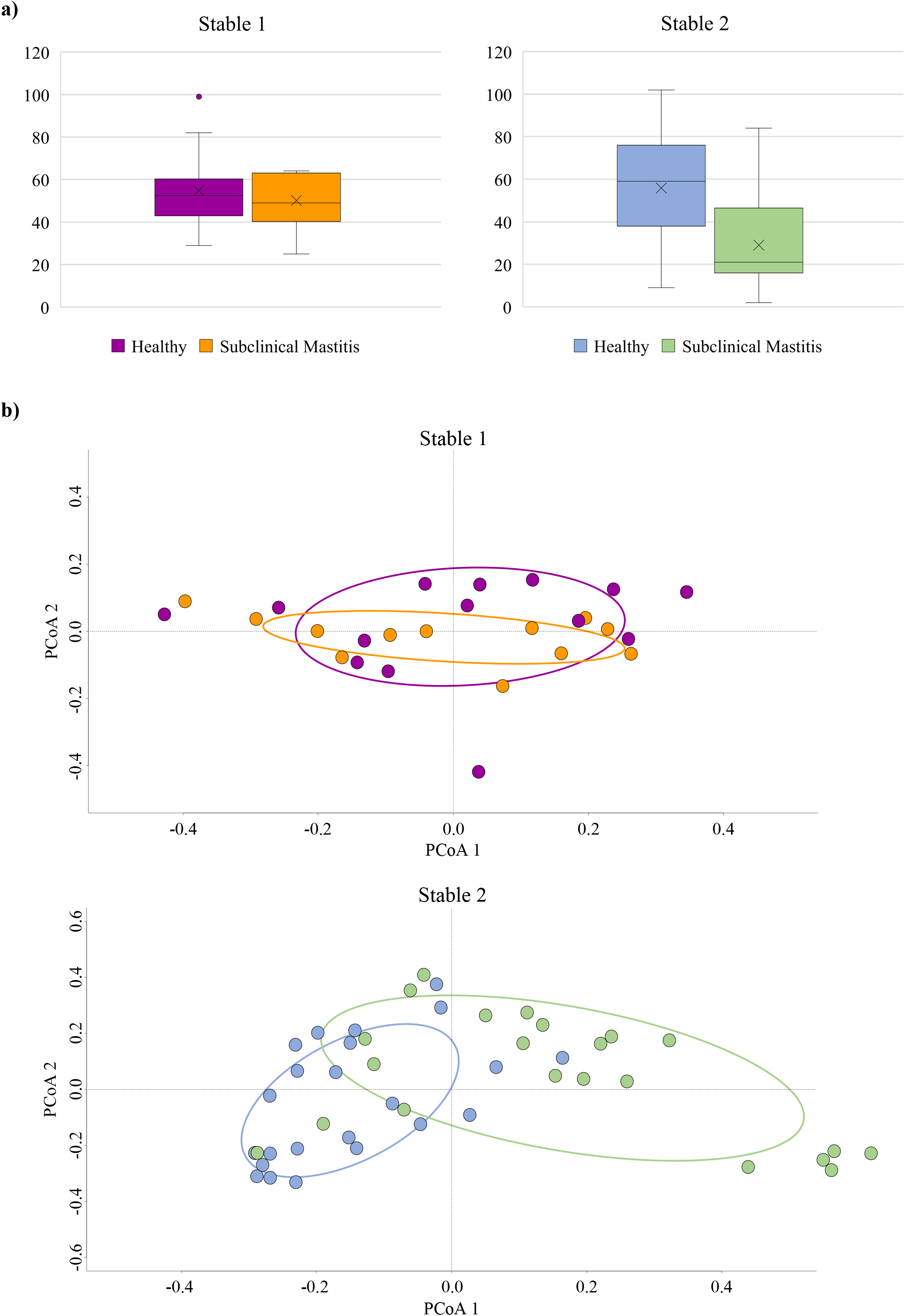
Species-level milk microbial biodiversity between healthy and SM-affected cows. Panel a shows the box and whisker plot of the calculated species-richness based on the number of microbial species observed between the two clinical status groups divided per stable. For each box and whisker plot, the x-axis reports the two considered clinical status-based groups, while the y-axis depicts the number of bacterial species. Boxes are determined by the 25^th^ and 75^th^ percentiles. The whiskers are determined by the maximum and minimum values that correspond to the box extreme values. Lines inside the boxers represent the average of the species number, while crosses correspond to the median. Panel b displays the two bidimensional Bray-Curtis dissimilarity index-based PCoA of each milk sample divided per stable.

### Subclinical mastitis effects on species-level core milk microbial communities

Reconstruction of the “core” milk microbiota, i.e., the bacterial taxa that are shared across samples of a defined cohort, allows the identification of the most prevalent bacterial species that inhabits the bovine milk (21, 22). In this context, to evaluate the impact that subclinical mastitis may have on the most prevalent milk bacterial species, the “core” microbial community characterizing milk samples from healthy cows was compared to that of milk from SM-affected bovines. Specifically, only those bacterial taxa with a prevalence > 80% were considered as part of the “core” milk community, as previously described (23). Based on this cut-off, 23 bacterial species resulted to be shared between the “core” microbiota of healthy and SM cows from stable 1, with *Aerococcus urinaeequi, Jeotgalibaca porci, Paraclostridium bifermentans, Romboutsia ilealis, Turicibacter sanguinis, Weissella jogaejeotgali* as well as two not yet identified species belonging to the genera *Romboutsia* and *Turicibacter* as the most abundant “core” taxa (average relative abundance >3%) (Table S4). Thus, suggesting that these microbial species are typical colonizers of the bovine milk regardless of the clinical status for stable 1. Conversely, three bacterial species, including *Bifidobacterium pseudolognum*, and two not yet characterized species belonging to the genera *Olsenella*, and *Staphylococcus* were exclusively part of the “core” microbiota of milk from healthy cows, while six microbial taxa only belonged to the “core” microbial community of subclinical mastitis milk samples, encompassing *Lactobacillus acidipiscis, Staphylococcus hominis*, and four unknown species of the genus *Anaerococcus, Jeotgalicoccus, Mogibacterium*, and *Tetragenococcus* (Table S4). Interestingly, *B. pseudolongum* has been identified as one of the main bifidobacterial players of the mammalian milk in healthy subjects (24-26), thus indicating that this bacterial species may be considered as marker of a healthy status that may undergo a reduction in prevalence in case of subclinical mastitis.

Differently from stable 1, in stable 2 only two bacterial species were shared between the “core” milk microbiota of healthy and SM-affected cows, i.e., two yet unclassified species belonging to the genera *Corynebacterium* and *Staphylococcus* (Table S4). Interestingly, the latter corresponded to the only two taxa present with a prevalence > 80% in the SM milk samples. In contrast, the “core” microbial community of milk collected from healthy cows consisted of 11 additional bacterial taxa including *Staphylococcus chromogenes, Clostridioides difficile*, and *R. ilealis* together with 8 not yet identified species belonging to genera *Clostridioides, Enterococcus, Kurthia, Lysinibacillus, Macrococcus, Paeniclostridium, Romboutsia, Staphylococcus*, and *Turicibacter* (Table S4). Thus, suggesting that, for stable 2, the inflammation of the mammary gland induced a more pronounced modulation of the “core” milk microbial community, when compared to that observed for stable 1, recording a drastic reduction in the number of bacterial species shared among the milk samples collected from cows with SM.

Overall, these results highlighted that SM does not induce standard modulation of the “core” milk bacterial composition, but rather, these microbial changes seem to depend on environmental factors. An observation that underlines the importance of characterizing the milk microbiota of healthy cows within each stable to create a “reference standard” to be compared with the microbial community of milk samples from subclinical mastitis-affected cows from the same stable to identify those microbial species potentially involved in the onset of SM for each stable. At the same time, the presence of yet unclassified species in the “core” microbiota of milk from both healthy and diseased cows highlighted the urgent need to apply culture-dependent approaches aimed at isolating and characterizing this milk microbial dark matter.

### Prediction of putative milk microbial markers correlated with subclinical mastitis

To identify possible microbial biomarkers strictly associated with SM, milk samples collected from healthy cows were compared with those from SM-affected cows. Interestingly, for stable 1, only one bacterial species, i.e., a not yet characterized species of the genus *Lactococcus*, was significantly more abundant in healthy cows when compared to the diseased ones (Student’s T-test p-value = 0.042), thus indicating that this taxon may be considered as a positive microbial biomarker associated with a healthy condition. However, the latter microbial species showed a reduced relative abundance (0.05%) as well as a low prevalence (26.67%) (Figure 2 and Table S4), thus suggesting that SM does not induce striking changes in the milk microbiota of stable 1, preventing the identification of SM-related microbial biomarkers. However, in depth insight into taxonomic profiles of milk samples from stable 1 highlighted that *Streptococcus dysgalactiae*, one of the most prevalent pathogens causing bovine mastitis worldwide, was detected only in milk samples from SM-affected cows, and when present, this species displayed a high average relative abundance (3.96%) (10, 27-29) (Table S4 and Table S5). At the same time, *Corynebacterium bovis*, another bacterial species listed among the pathogenic microorganisms closely associated with SM, possessed a higher average relative abundance and prevalence in milk samples from cows with SM with respect to the ones collected from healthy cows (Table S4) (30-32). In this context, even if not statistically significant, these results strengthen the notion that both *S. dysgalactiae* and *C. bovis* may be considered as microbial biomarkers of SM.

**Figure 2:**
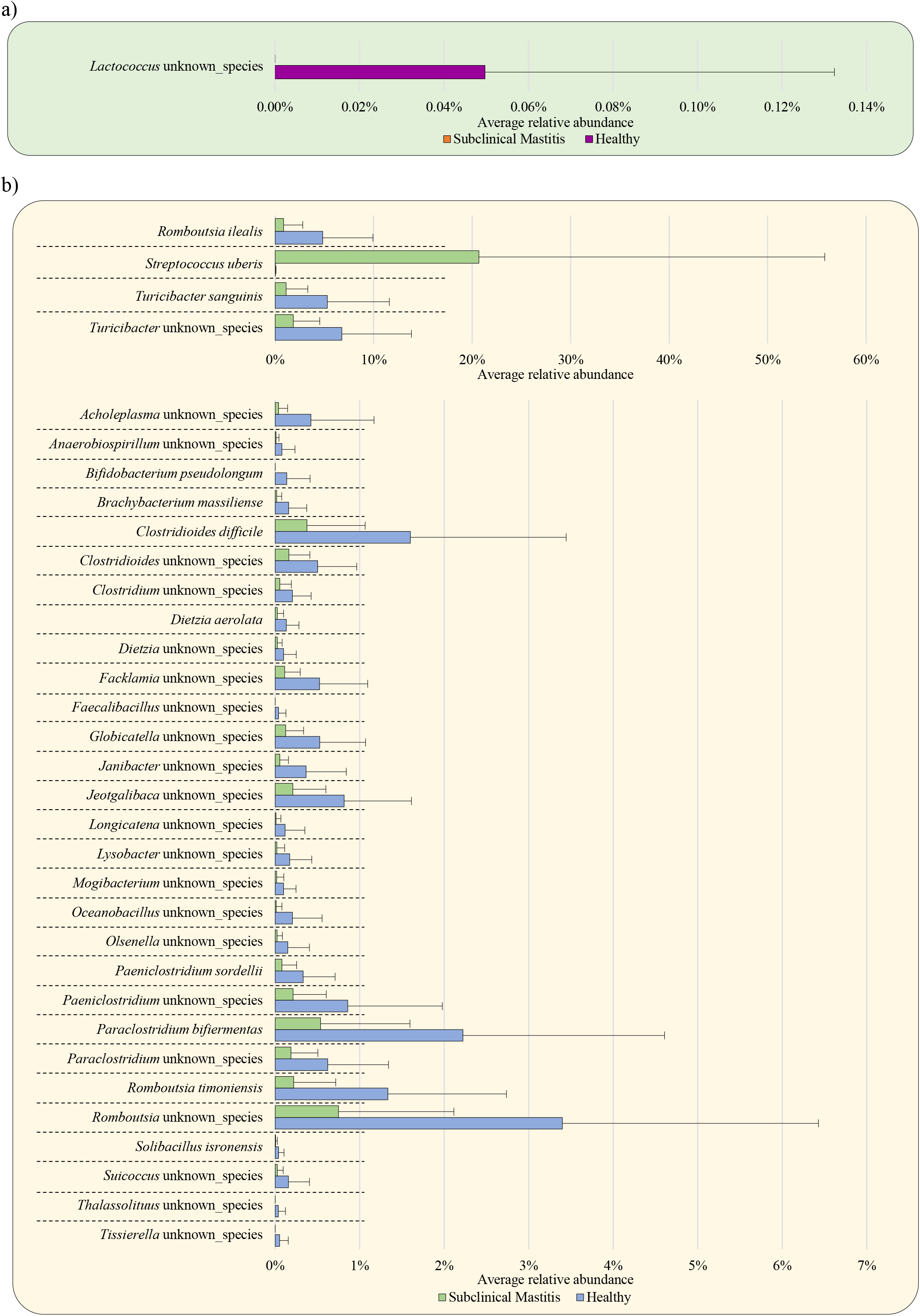
Species-level differences in the taxonomic composition of milk samples from healthy and SM-affected cows. Panels a and b report the average relative abundances of those bacterial species that significantly differ in the milk samples from healthy and diseases cows for stable 1 and 2, respectively.

Differently from stable 1, the comparison of taxonomic profiles between healthy and diseased cows from stable 2 highlighted that the average relative abundance of 33 bacterial species significantly differed based on the clinical status (Figure 2 and Table S5). Among the latter, *B. pseudolongum* was only found in milk samples from healthy cows (Student’s T-test *p*-value = 0.025) (Figure 2 and Table S5). In this context, as above reported, since *B. pseudolongum* has been identified as a commensal microorganisms of bovine milk and members of the genus *Bifidobacterium* are known to play multiple beneficial effects upon their host promoting anti-inflammatory response, providing protection against pathogen colonization, and favoring the proliferation of beneficial butyrogenic microbial players that can use the acetate produced by the bifidobacterial fermentation of complex glycans, this species can be considered as microbial biomarker of a healthy status (7, 30, 33-35). Furthermore, *Dietzia aerolata*, as well as three yet unclassified species of the genera *Dietzia, Facklamia*, and *Janibacter* were not only more prevalent but also significantly more abundant in milk samples from healthy cows when compared to that of subjects with SM (Figure 3, Table S4 and Table S5). Notably, these genera were considered as commensal microorganisms of the milk microbiota (36-38) suggesting their possible involvement as positive microbial markers of a healthy conditions. However, the fact that these microbial species corresponded to taxa not yet isolated strengthen the notion that a culture-based research effort is required for the isolation and characterization of potential microbial markers of a healthy or SM status. Conversely, *Streptococcus uberis* was identified as the only bacterial species, among those taxa that significantly differed between the two cow groups, with a significantly higher relative abundance in the milk samples of cows with SM when compared to the those from healthy cows (Student’s T-test *p*-value = 0.013) (Figure 3, Table S4 and Table S5). *S. uberis* has been widely described as a common pathogen strictly related with both clinical and subclinical mastitis thanks to its ability to persist under environmental stress or exposure to antibiotic treatment inducing biofilm formation when in contact with α- and β-casein milk component (39-42). Thus, indicating *S. uberis* as the potential responsible microorganism of the bovine intramammary infection in stable 2 and electing this taxon as biomarker of subclinical mastitis.

**Figure 3:**
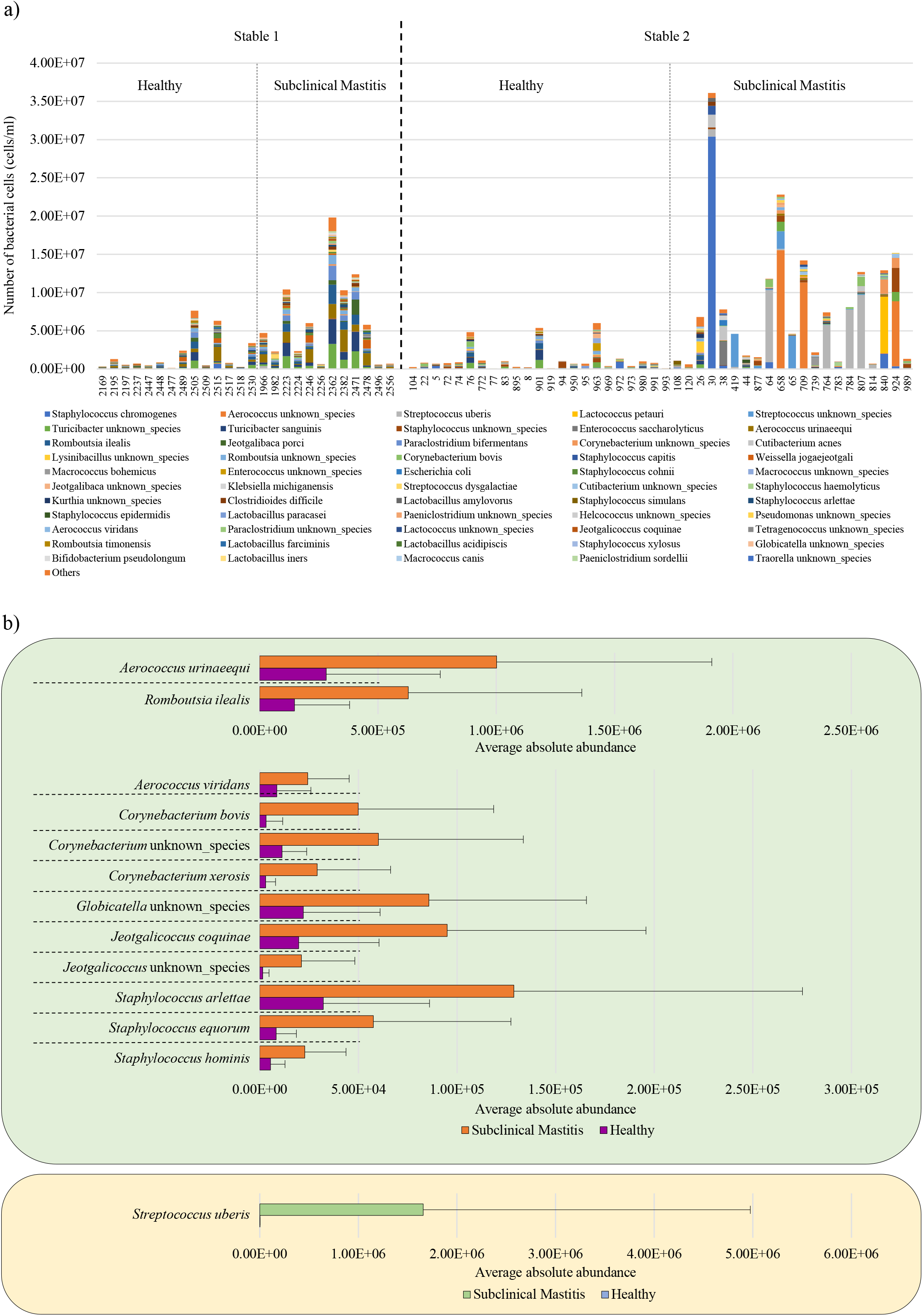
Evaluation of milk total bacterial counts between healthy and SM-affected cows. Panel a reports the bar plot showing the taxonomic profiles of each milk sample normalized for the flow cytometry-based bacterial cell enumeration. Panel b and c display the microbial species whose absolute abundance differed between healthy and SM-affected cows in stable 1 and 2, respectively.

Overall, these results not only reinforce the above-mentioned evidence that SM induces different variations in the microbial composition of bovine milk depending on the environment, but also underline that the microbial etiological agents of SM differ across stables. Thus, suggesting that the characterization and identification of the microbial agents causing SM is essential to activate targeted strategies to limit the diffusion of the intramammary gland infection.

### Bacterial cell count-dependent taxonomical differences in the milk microbiota of healthy and subclinical mastitis-affected cows

To evaluate whether SM may have an impact on the overall number of the bacterial cells present in bovine milk, each collected milk sample was subjected to a flow cytometry-based total bacterial cell enumeration. Interestingly, for both stables, the average of the microbial cell number present in the milk samples collected from cows with SM was significantly higher when compared to that observed for samples from healthy cows (Student’s T-test p-value of 0.002 and 0.001 for stable 1 and 2, respectively) (Figure 3 and Table S6). Indeed, the average of the flow cytometry readouts related to milk samples from cows with SM exceeded by at least 3 times the observed average number of bacterial cells in milk samples from healthy cows for both stables, with an average of 1.84E+06 and 1.26E+06 cells/ml for milk from healthy cows and 6.33E+06 and 8.34E+06 cells/ml for milk from SM-affected cows for stable 1 and 2, respectively (Figure 3 and Table S6). An observation that leads to suggest that this inflammatory disease affecting the bovine mammary glands is not only characterized by an alteration of the milk microbial community, but also by a significant increase in the number of microbial cells present in the milk. Furthermore, differently from milk taxonomic composition that undergoes different alterations among stables, an increase in the number of bacterial cells in SM cow-derived milk samples seemed to be a common feature of the two different stables regardless of environmental factors.

Based on these observations, to obtain a comprehensive biological interpretation of the analyzed milk microbial community complexity and to identify further differences in the taxonomic composition of milk samples between healthy and SM-affected cows based on the number of bacterial cells, the assessed cell counts were subsequently employed to normalize ITS microbial profiling sequencing data transforming relative metagenomic data into absolute abundances, as previously described (20). Insights into the latter revealed that the number of cells of 12 bacterial species significantly differed between milk samples from healthy and SM-affected cows in stable 1 (Figure 3). Interestingly, *C. bovis*, whose relative abundance was not significant between the two groups as above reported, displayed a significantly higher absolute abundance in milk samples from cows with mammary gland inflammation (average absolute abundance of 4.99E+04 cells/ml) when compared to the healthy ones (average absolute abundance of 3.14E+03 cells/ml) (Student’s T-test p-value = 0.038) (Figure 3 and Table S6). Furthermore, *Corynebacterium xerosis*, another bacterial species frequently associated with bovine subclinical mastitis (43-45), showed a significant average absolute abundance increment of almost 10 times, moving from 2.94E+03 to 2.91E+04 cells/ml in milk samples from cows with intramammary infection when compared to the healthy ones (Student’s T-test p-value = 0.033) (Figure 3 and Table S6). In this context, the evaluation of the absolute abundances allowed to identify two species of the genus *Corynebacterium*, i.e., *C. bovis* and *C. xerosis*, as the potential etiological agents of SM for stable 1. Moreover, since the two bacterial species are not exclusively present in the microbial community of milk from cows with inflammation of the mammary gland, it is possible to assume that a certain cell number of these two species is necessary to induce the inflammatory condition typical of SM.

Conversely, the assessment of the absolute abundance-based taxonomic profiles for stable 2 revealed that only a single species significantly differed between milk samples of healthy and diseased cows, i.e., *S. uberis* (Student’s T-test *p*-value = 0.033) (Figure 3 and Table S6). Specifically, this microbial species displayed an increment of the cell number of almost 5-fold in the milk samples from cows with SM when compared to that of the healthy cows (Figure 3 and Table S6). Thus, confirming the role of this species in the onset of subclinical mastitis in stable 2. Overall, these results highlighted how the comparison of absolute abundances, obtained through the combination of a sequencing approach with a flow cytometry-based total cell count, may provide more accurate information about the alteration that the milk microbial composition may undergo in case of SM.

## Conclusions

Bovine intramammary inflammation represents a worldwide burden causing serious repercussions not only on the health of dairy herds, but also on milk productivity and quality (15, 46). To limit the spread of this disease, and especially of its silent form, i.e., subclinical mastitis, whose containment is difficult due to the lack of evident symptoms and its high incidence rate, the identification of SM microbial causative agents is of crucial significance (12, 47-49). However, the impact of bovine milk microbial composition that may have on SM has not yet been fully investigated. In this context, the application of an ITS microbial profiling to milk samples collected from healthy and SM-affected cows from two different stables highlighted that environmental factors play a predominant role in the modulation of the milk microbial community regardless of the clinical status, thus suggesting the need to separately analyze samples according to their stable of origin to avoid environmental factor-related biases. The subsequent comparison of milk samples divided per stable showed that, in general, SM is associated to a reduced number of species in the milk microbial community. On the contrary, the analyses of the impact that SM may have on the “core” milk microbiota between healthy and diseased bovines showed that this silent intramammary inflammation induces stable-related alteration of the “core” milk bacterial composition. Thus, emphasizing that, despite being characterized by a lack of obvious symptoms, subclinical mastitis is accompanied by an alteration of the bovine milk microbiota. Furthermore, the enumeration of the bacterial cells presents in each collected milk sample through flow cytometry evidenced that, in general, SM is associated with a significant increase of milk bacterial cells. Furthermore, the normalization of the metagenomic taxonomic profiles with the obtained total cell counts allowed to obtain absolute abundance-based compositional profiles that not only identified *Corynebacterium bovis* together with *Corynebacterium xerosis* and *Streptococcus uberis* as potential microbial markers of SM for stable 1 and 2, respectively, but also lead to suggest that the intramammary inflammation typical of SM may not only be associated with the presence of a certain bacterial taxon, but also with the total number of cells of that species in the bovine milk. Thus, suggesting the importance of the combination of a sequencing approach with a bacterial cell enumeration to obtain a more accurate overview of the milk microbial composition associated with subclinical mastitis.

## Acknowledgements

We thank GenProbio srl for financial support of the Laboratory of Probiogenomics. Part of this research has been conducted using the High-Performance Computing (HPC) facility of the University of Parma.

## Supplementary figure legend

**Figure S1:**
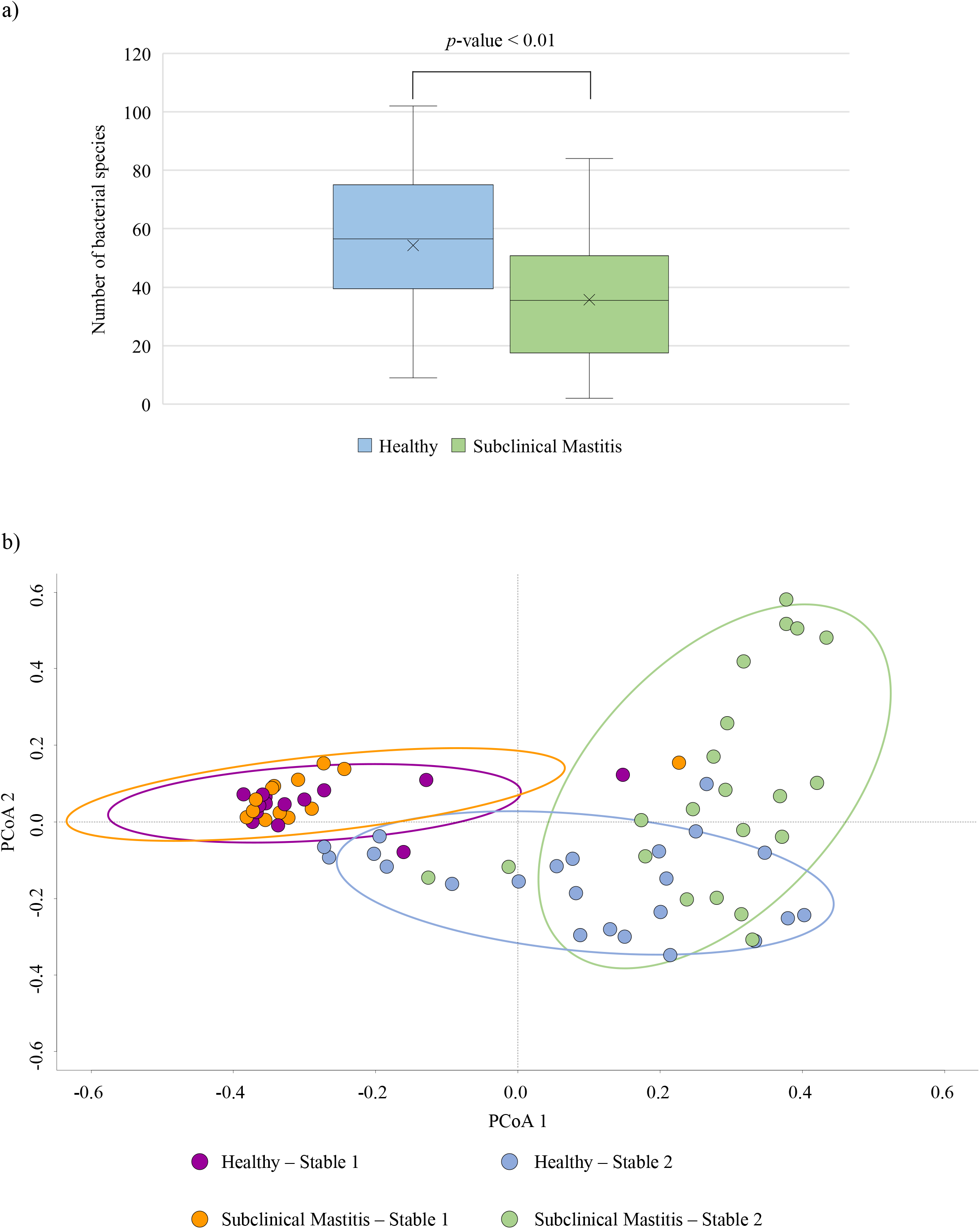
Differences in milk microbial biodiversity between healthy and SM-affected dairy cows. Panel a shows the box and whisker plot of the calculated species-richness based on the number of microbial species observed between the two clinical status groups. For each box and whisker plot, the x-axis reports the two considered clinical status-based groups, while the y-axis depicts the number of bacterial species. Boxes are determined by the 25^th^ and 75^th^ percentiles. The whiskers are determined by the maximum and minimum values that correspond to the box extreme values. Lines inside the boxers represent the average of the species number, while crosses correspond to the median. Panel b displays the two bidimensional Bray-Curtis dissimilarity index-based PCoA of each collected milk sample. The ellipses of the PCoA were drawn based on the standard deviation of each considered group.

